# Molecular mechanism of the endothelin receptor type B interactions with Gs, Gi, and Gq

**DOI:** 10.1101/2024.04.29.591572

**Authors:** Donghee Ham, Wataru Shihoya, Osamu Nureki, Asuka Inoue, Ka Young Chung

**Affiliations:** School of Pharmacy, Sungkyunkwan University, 2066 Seobu-ro, Jangan-gu, Suwon 16419, Republic of Korea; Department of Biological Sciences, Graduate School of Science, The University of Tokyo, 7-3-1 Hongo, Bumkyo-ku, Tokyo, 113-0033, Japan; Graduate School of Pharmaceutical Sciences, Tohoku University, 6-3, Aoba, Aramaki, Aoba-ku, Sendai, Miyagi, 980-8578, Japan

**Keywords:** ET_B_, G protein, Signal selectivity, HDX-MS, Conformation

## Abstract

The endothelin receptor type B (ET_B_) exhibits promiscuous coupling with various G protein subtypes including Gs, Gi/o, Gq/11, and G12/13, leading to multi-modal signal transduction. Recent fluorescence and structural studies have raised questions regarding the coupling efficiencies and determinants of these G protein subtypes. Herein, by utilizing an integrative approach, combining hydrogen/deuterium exchange mass spectrometry and NanoLuc Binary Technology-based cellular systems, we investigated conformational changes of Gs, Gi and Gq triggered by ET_B_ activation. ET_B_ coupled to Gi and Gq but not with Gs. We underscored the critical roles of specific regions, including the C-terminus of Gα and intracellular loop 2 (ICL2) of ET_B_ in ET_B_-Gi1 or ET_B_-Gq coupling. Although The C-terminus of Gα is essential for ET_B_-Gi1 and ET_B_-Gq coupling, ET_B_ ICL2 influences Gq-coupling but not Gi1-coupling. Our results suggest a differential coupling efficiency of ET_B_ with Gs, Gi1, and Gq, accompanied by distinct conformational changes in G proteins upon ET_B_-induced activation.

## INTRODUCTION

Endothelin receptors are class A G protein-coupled receptors (GPCRs) that are abundantly expressed in endothelial and vascular smooth muscle cells and regulate numerous cellular signaling pathways, including vascular, renal, pulmonary, coronary, and cerebral circulation^1^. There are two subtypes of endothelin receptors: endothelin receptor type A (ET_A_) and B (ET_B_)^2,3^. Both ET_A_ and ET_B_ mediate phospholipase (PL) A2, PLC, and PLD activation through diverse G protein couplings, including Gq/11, Gs, and Gi/o^4^. ET_A_ has been studied primarily for their Gs coupling, leading to adenylyl cyclase signal transduction. However, it remains unclear whether ET_B_ mediates activation of adenylyl cyclases directly via Gs coupling or indirectly via PLC activation^4,5^. Consequently, delving into the selectivity and promiscuity of G protein coupling is indispensable for understanding ET_B_-mediated G protein signaling.

Efforts have been made to label the magnitude of ET_B_-G protein coupling^6–9^. The International Union of Basic and Clinical Pharmacology and the British Pharmacological Society Guide to Pharmacology (GtP), an expert-curated literature database, have outlined “primary coupling” as the principal coupling mode and “secondary coupling” as its complementary counterpart, albeit without quantitative metrics^10^. Currently, quantitative datasets are accessible through GPCRdb (GPCRdb.org), encompassing assays such as the transforming growth factor-α (TGF-α) shedding assay and the G protein effector membrane translocation assay^11^. ET_B_ is curated as a promiscuous receptor that has similar coupling magnitude throughout G protein subtypes by GtP and TGF-α shedding assay^7,10^. However, a recently published BRET-based free βγ assay suggested that ET_B_ couples to Gi/o primarily and does not couple to Gs^9^.

Recent cryo-EM studies of ET_B_ in complex with Gq or Gi1 have provided insight into coupling mechanisms and G protein-subtype selectivity between ET_B_ and G proteins^12–14^. In these studies, however, owing to weak or unstable association between ET_B_ and G proteins, the ET_B_-G protein complexes were forced to be made using a HiBiT tethering strategy^15^. Moreover, the G proteins were heavily modified; Gαq is engineered based on mini-Gαs, and αN is replaced by Gαi. Investigating the mechanism of G protein coupling in the ET_B_ using un-modified proteins could offer vital understanding of innate interactions and operational mechanisms, which has yet been explored.

Herein, we studied the coupling selectivity and conformational dynamics in un-modified or mimimally modified ET_B_ and G proteins using an integrative approach involving hydrogen/deuterium exchange mass spectrometry (HDX-MS) and NanoLuc Binary Technology (NanoBiT)-based cellular biosensor systems. By comparing Gs, Gi1, and Gq, we elucidated the roles of specific regions in their coupling mechanisms.

## RESULTS AND DISCUSSION

### Differential coupling efficiency of ET_B_ with Gs, Gi1, and Gq within the cell

Although ET_B_ is recognized to activate Gs, Gi1, and Gq subfamilies (GPCRdb.org), the relative coupling efficiencies have not been examined. We performed a NanoBiT-based G protein assay to compare the coupling efficiencies of ET_B_-Gs, ET_B_-Gi1, and ET_B_-Gq^7,15^. To monitor the G protein activation, Gα was fused with the NanoBiT large fragment (LgBiT) and Gγ_2_ was fused with the small fragment (SmBiT) at the N-terminus (Figure 1A). Upon GPCR-induced G protein activation, Gα undergoes GDP/GTP turnover and GTP-bound Gα dissociates from Gβγ, leading to the dissociation of LgBiT and SmBiT, thereby decreasing luminescent signals. The NanoBiT-G protein assay showed that the extent of ET_B_-induced G protein activation varied among G protein subtypes (Figure 1B). Gi1 and Gq exhibited the significant activation, while Gs showed minimal activation. This result is consistent with a recent report, which used free βγ assay^9^.

**Figure 1.**
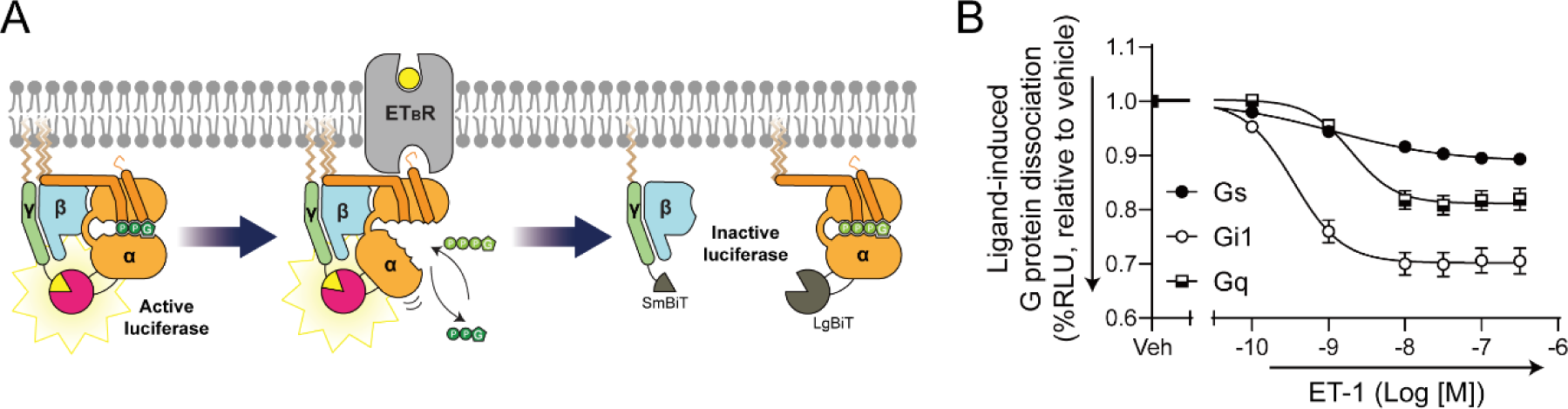
Differential efficiency of ET_B_-mediated G protein activation. (A) Schematic representation of the NanoBiT-based Gα-Gβγ dissociation assay. ET_B_ was expressed together with the Gαs–LgBiT or Gαi1–LgBiT or Gαq–LgBiT, Gβ1, and Gγ2-SmBiT constructs. (B) ET_B_-mediated G protein activation. ET1-induced Gα and Gβγ dissociation was measured using the NanoBiT assay. Symbols and error bars represent mean and standard error of the mean, respectively, derived from three independent experiments, each conducted in duplicate.

### Differential conformational changes of Gi1, Gq, and Gs upon co-incubation with ET_B_

To compare the dynamic conformational changes in G proteins upon ET_B_-binding, HDX-MS was performed. HDX-MS measures the exchange rate between the amide hydrogen atoms in the proteins and deuterium in the solvent, revealing the conformational dynamics of the proteins^16,17^. This technique has been successfully used to study conformational changes upon protein-protein interactions, even with weak or unstable association^18,19^. Therefore, HDX-MS is an appropriate technique for studying the conformational changes induced by weak ET_B_-G protein interactions.

We prepared GDP-bound heterotrimeric Gs, Gi1, and Gq, along with ET-1-bound ET_B_. The G proteins were incubated with or without ET-1-bound ET_B_, and then HDX was induced by incubating in the D_2_O buffer for 10, 100, 1,000, or 10,000 s. The proteins were fragmented by pepsin digestion, and the detailed information about the peptic peptides used for HDX-MS analyses are described in Figure S1 and Supplementary dataset.

The Gαi1 and Gαq, co-incubation with ET_B_, showed higher HDX near the nucleotide-binding pocket (*i.e.*, p-loop through α1, αE/αF loop through αF, β5 through αG, and β6/α5 loop) than those of Gαi1 and Gαq without co-incubation with ET_B_ (Figure 2, Supplementary dataset). Higher HDX levels near the nucleotide-binding pocket implied that these regions were exposed to the buffer and/or became more dynamic. In our previous GPCR-G protein coupling studies, these HDX profiles indicated GDP release^19–21^. Therefore, these results suggest that ET_B_ co-incubation induced GDP release from Gi1 and Gq, indicating functional coupling between purified ET_B_ and Gi1/Gq. Unlike Gi1 and Gq, the HDX profiles of Gs were not altered upon co-incubation with ET_B_ (Supplementary dataset), indicating that purified ET_B_ did not efficiently couple to Gs, which is consistent with the NanoBiT assay data (Figure 1B).

**Figure 2.**
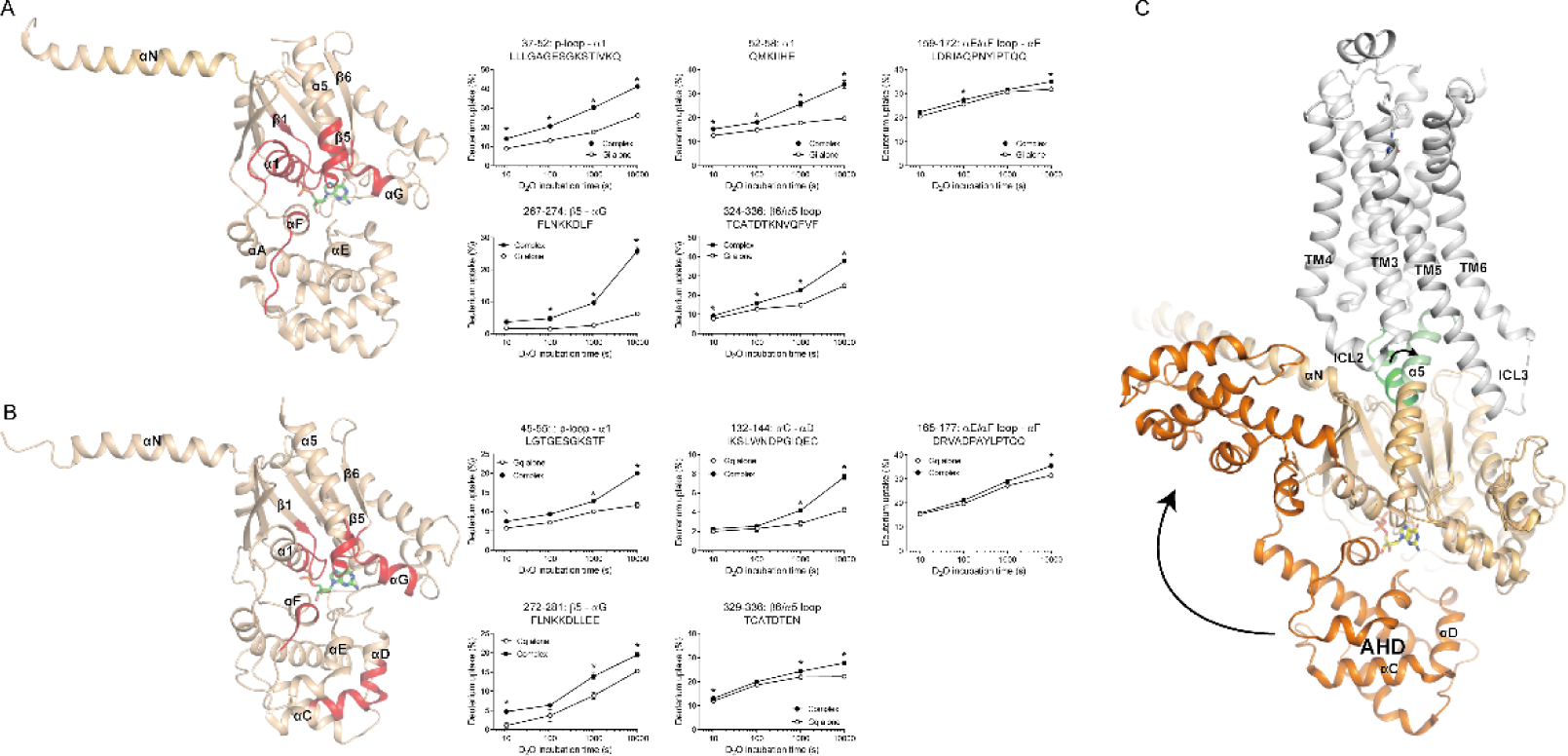
HDX profiles of Gi1 and Gq upon ET_B_ co-incubation. (A) Changes in the HDX profile of Gαi1 upon co-incubation with ET-1-bound ET_B_. The regions exhibiting increased HDX are colored red on the crystal structure of GDP-bound inactive Gi1 (PDB: 1GP2), with corresponding deuterium uptake plots for selected peptides presented as graphs. (B) Changes in the HDX profile of Gαq upon co-incubation with ET-1-bound ET_B_. The regions exhibiting increased HDX are colored red on the crystal structure of GDP- and YM-254890-bound inactive Gαq (PDB: 3AH8), with corresponding deuterium uptake plots for selected peptides presented as graphs. The results in (A) and (B) were obtained from three independent experiments, and the statistical significance of differences was assessed using Student’s t-test (*p <0.05). Data are represented as mean ± standard error of the mean. (C) Superposition of structures of β_2_AR-Gs complex (PDB: 3SN6) and GDP-bound Gs (PDB: 6EG8). β_2_AR is colored grey. Ras-like domain of Gαs is colored light orange, α- helical domain of Gαs is colored orange, and C-terminus of Gαs is colored green.

Intriguingly, αC through αD of Gαq showed higher HDX levels in Gq co-incubated with ET_B_ than in Gq alone (Figure 2B, peptide 132-144). This region is located at the α- helical domain (AHD) of Gα (Figure 2C, orange). Although the functional significance of AHD in G protein signaling remains poorly studied, it has been well-established that the AHD is displaced in the nucleotide-free Gα (Figure 2C)^22^. Therefore, the higher HDX levels in αC through αD in Gq incubated with ET_B_ than in Gq alone may reflect the increased local conformational dynamics within the AHD due to AHD displacement upon GDP release. However, we did not detect altered HDX levels within the Gαi AHD upon ET_B_ co-incubation (Figure 2A). Therefore, the increased local dynamics within the AHD is likely dependent on Gα subtypes. Previously, we observed higher HDX in the local region of the AHD in the M3-Gq^19^ and β_2_AR-Gs^20,23^ studies but not in the M2-Gi/o study^21^. As the increased local conformational dynamics was also detected within the AHD in the current ET_B_-Gq (Figure 2B) complex but not in the current ET_B_-Gi1 (Figure 2A), it is speculated that, upon GDP release, the local conformational dynamics of Gα AHD is affected in Gαs or Gαq only.

### Binding interfaces between Gi1/Gq and ET_B_

We expected that the HDX levels would decrease at the ET_B_ and G protein interfaces upon co-incubation of ET_B_ and G protein because the amide hydrogens would be protected from the buffer at the protein-protein interaction interfaces. Indeed, in our previous HDX-MS studies, the C-terminal part of α5 (Figure 2C, green) of Gαs, Gαo/i, and Gαq showed reduced HDX levels when their primary upstream GPCRs were co-incubated^20,21,24^ because this region is the most-well conserved receptor-binding interface. However, no regions of Gαi1 and Gαq showed reduced HDX levels upon ET_B_ co-incubation (Figure 2A and 2B, Supplementary dataset). Yet, as described above, the HDX profile showed higher HDX levels at the nucleotide-binding regions in the ET_B_-co-incubated Gαi1 or Gαq (Figure 2A and 2B), supporting ET_B_-induced GDP release from Gi1 and Gq in the current HDX system. Together, the data suggests the shallow or unstable nature of the ET_B_-Gi1 or ET_B_-Gq interaction and explain the necessity for HiBiT tethering strategies for stable ET_B_-Gi1 or ET_B_-Gq complex formation^12,13^.

We also expected that the intracellular parts of ET_B_ would reveal lower HDX levels after co-incubation with G proteins as intracellular components such as intracellular loop 2 (ICL2), ICL3, cytosolic regions of transmembrane domain 5 (TM5), or TM6 interact extensively with G proteins (Figure 2C). In the ET_B_ side, HDX signals were not detected at most of the cytosolic regions except for helix 8 and the C-tail (Figure 3A, Supplementary dataset), due to the absence of the mass spectrum of the peptic peptides in these regions (Figure S1C). Since the cytosolic regions of the receptor are the binding interfaces for G proteins, information regarding the binding interfaces, with the current ET_B_-G protein HDX-MS studies, could not be obtained.

**Figure 3.**
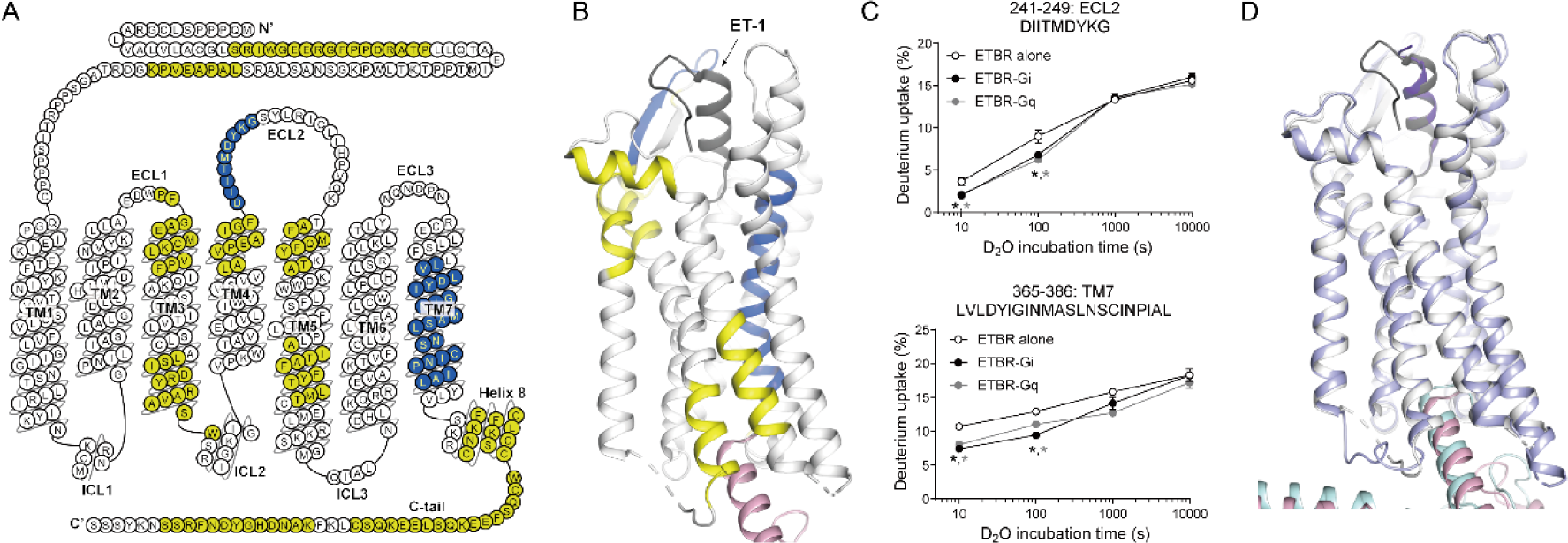
HDX profiles of ET_B_ upon Gq or Gi1 co-incubation. (A and B) Changes in the HDX profile of ET-1-bound ET_B_ upon co-incubation with heterotrimeric Gq. The regions exhibiting decreased HDX are colored blue in the ET_B_ snake plot (A) and on the crystal structure of ET-1-bound active ET_B_ (PDB: 8HCX) (B). The regions where HDX levels were not changed are colored yellow, and the regions where peptides were not identified are colored white. (C) Deuterium uptake plots of selected peptides from the color-coded regions of ET_B_ are shown as graphs. Results were obtained from three independent experiments, and the statistical significance of differences was assessed using Student’s t-test (*p <0.05). Data are represented as mean ± standard error of the mean. (D) Superposition of structures of ET_B_-Gi1 complex (PDB: 8HBD) and ET_B_-Gq complex (PDB: 8HCX). Gi1-coupled ET_B_ is colored light purple and Gq-coupled ET_B_ is colored white. Gαi1 is colored light cyan and Gαq is colored light pink.

### Conformational changes of ET_B_ upon co-incubation with Gs, Gi1, or Gq

Owing to the nature of the transmembrane protein, the sequence coverage of the mass spectrometric analysis of ET_B_ was low (41%) (Figure S1C and Figure 3A). Nevertheless, with the analyzed peptic peptides, we observed that the HDX-MS profiles of Gi1 and Gq co-incubated ET_B_ were identical (Supplementary dataset), consistent with the similar high-resolution structures of ET_B_ complexed with Gi1 and Gq (Figure 3D)^12,13^.

Upon interaction with Gi1 or Gq, ET_B_ showed reduced HDX levels within the extracellular loop 2 (ECL2) and TM7 (Figure 3A–3C, Supplementary dataset), suggesting that these regions became more rigid or less exposed to the buffer upon co-incubation with Gi1 or Gq. ECL2 is at the extracellular side, therefore, the conformational change at ECL2 would be due to allosteric conformational changes transmitted from the G protein binding sites at the intracellular regions. This might be explained by a previous study that reported the contraction of the ligand-binding pocket upon G protein binding^25^. The changes in the HDX levels at TM7 may reflect the movement of TM regions upon G protein binding.

### Role of the C-terminus of Gα α5 in ET_B_-Gi1 or ET_B_-Gq coupling

The C-terminus of Gα α5 is the major binding interface of GPCR-G protein complexes (Figure 2C, green). This region is widely recognized to be critical for mediating interactions with receptors as well as for determining selective coupling between a receptor and G protein ^26–28^. Indeed, our previous HDX-MS studies showed decreased HDX levels in this region upon co-incubation with receptors, and truncation of the last five residues from the C-terminus of Gα α5 (Gα_Δ5) impeded primary Gs-, primary Gi/o-, and primary Gq-couplings (i.e., β_2_AR-Gs, M2-Gi/o, and M3-Gq, respectively)^21,24^. Curiously, in the secondary coupling (*i.e.*, β_2_AR-Gi coupling), the HDX levels of the C-terminus of Gαi/o remained unchanged upon co-incubation with β_2_AR, and Gαi_Δ5 minimally affected the β_2_AR-induced GDP release^21^, suggesting varying contributions of the C-terminus of Gα α5 to the coupling process in GPCR-G protein pair-dependent manners.

The high-resolution structures of ET_B_-Gi1 and ET_B_-Gq complexes also revealed extensive interaction of the C-terminus of Gα α5 with the receptor (Figure 4A and 4B)^12,13^. However, we did not observe HDX changes at the C-terminus of Gα upon co-incubation with ET_B_ (Figure 2A and 2B, Supplementary dataset). The discrepancy between the cryo-EM structures and HDX-MS study may be due to the different experimental systems (*i.e.*, HiBiT-aided stable complex formation between ET_B_ and miniG in the cryo-EM studies *vs*. un-modified proteins in the current HDX-MS studies). Therefore, further investigation of the involvement of the C-terminus of Gα α5 in ET_B_-G protein coupling is necessary.

**Figure 4.**
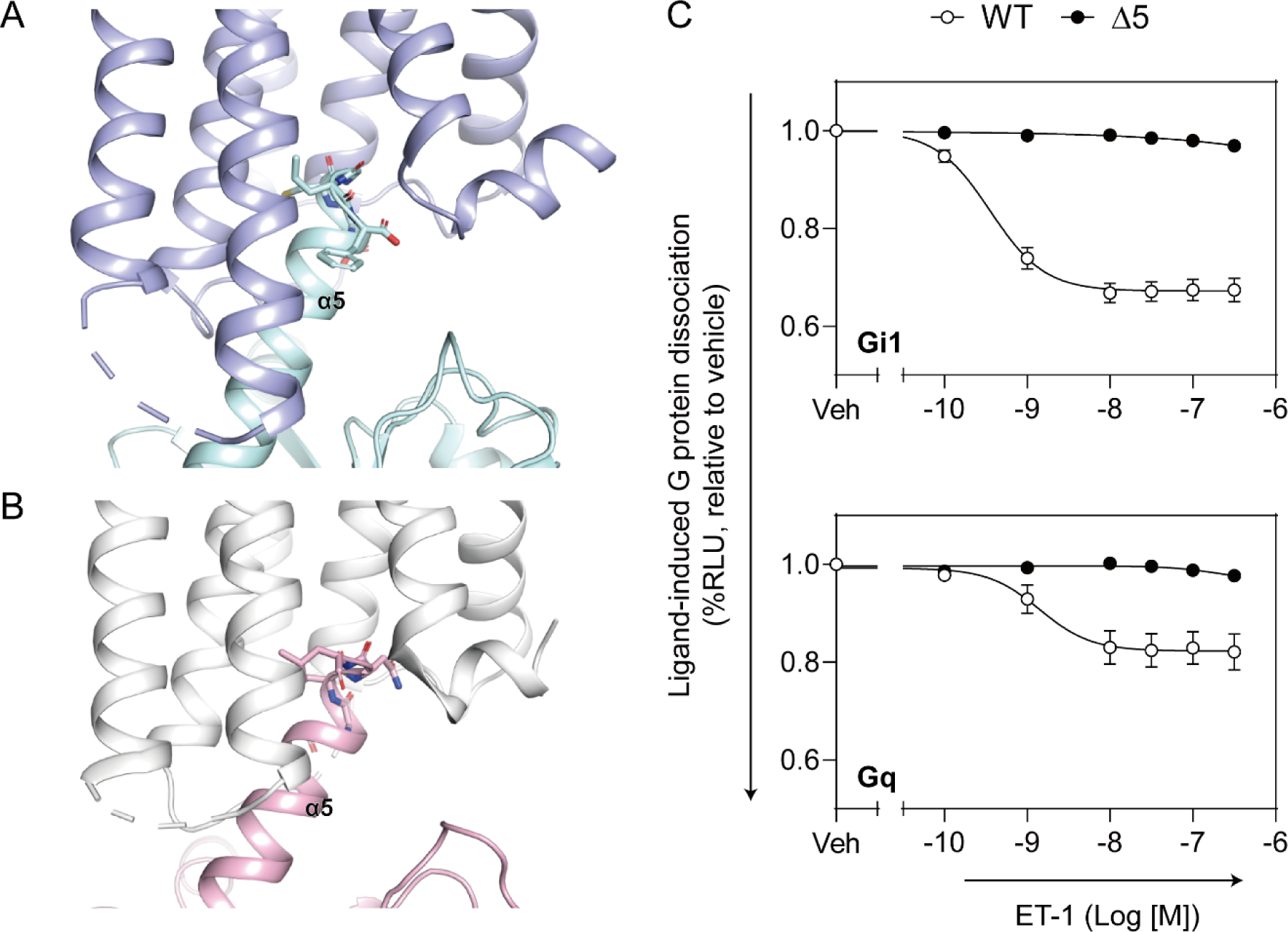
Role of Gα α5 on ET_B_-mediated G protein activation. (A and B) The interaction of ET_B_ with Gαi1 α5 (A) and ET_B_ with Gαq α5 (B). Selected residues are represented as sticks on the crystal structures of ET_B_-Gi1 (PDB: 8HBD) and ET_B_-Gq (PDB: 8HCX). The receptors are colored light purple and white, Gαi1 is colored light cyan, and Gαq is colored light pink. (C) Effects of Gα C-terminal deletion on ET_B_-mediated G protein activation. HEK293 cells expressing ET_B_ together with the WT or Δ5 Gα–LgBiT, Gβ1, and Gγ2-SmBiT constructs were subjected to the NanoBiT-G protein assay. Symbols and error bars represent mean and standard error of the mean, respectively, derived from three independent experiments, each conducted in duplicate. In many data points in the Δ5 constructs, error bars are smaller than the size of the symbols.

To examine the importance of the C-terminus of Gα α5 in coupling with ET_B_, we performed a functional study using mutant constructs. We generated C-terminal truncated Gα_Δ5 and analyzed ET_B_-induced G protein activation using the NanoBiT-G protein assay (Figure 1A). We assessed the Gα_Δ5 constructs for Gαi1 and Gαq, both of which showed relatively potent dissociation signals (Figure 1B) and found that ET_B_ failed to activate Gαi1_Δ5 and Gαq_Δ5 (Figure 4C). This suggests that the C-terminus of Gα α5 is still critical for ET_B_-Gi1 and ET_B_-Gq coupling, although HDX-MS data suggests a shallow or unstable interaction between ET_B_ and the C-terminus of Gαi1 or Gαq.

### Role of ICL2 in ET_B_-Gi1 or ET_B_-Gq coupling

Besides the C-terminus of Gα α5, ICL2 of a GPCR is well-studied for its role in the GPCR-mediated GDP release from a G protein and determination of the GPCR-G protein coupling selectivity^26^. Specifically, the residue 34.51 in ICL2 (residue numbering is based on the GPCRdb numbering scheme^29^) is mostly conserved as a bulky hydrophobic residue (Figure 5A), enabling extensive hydrophobic contacts with the Gα hydrophobic pocket in GPCR-Gs or GPCR-Gq complexes (Figure S2A and S2B), however this interaction is relatively shallow in the GPCR-Gi/o complexes (Figure S2C)^30–32^.

**Figure 5.**
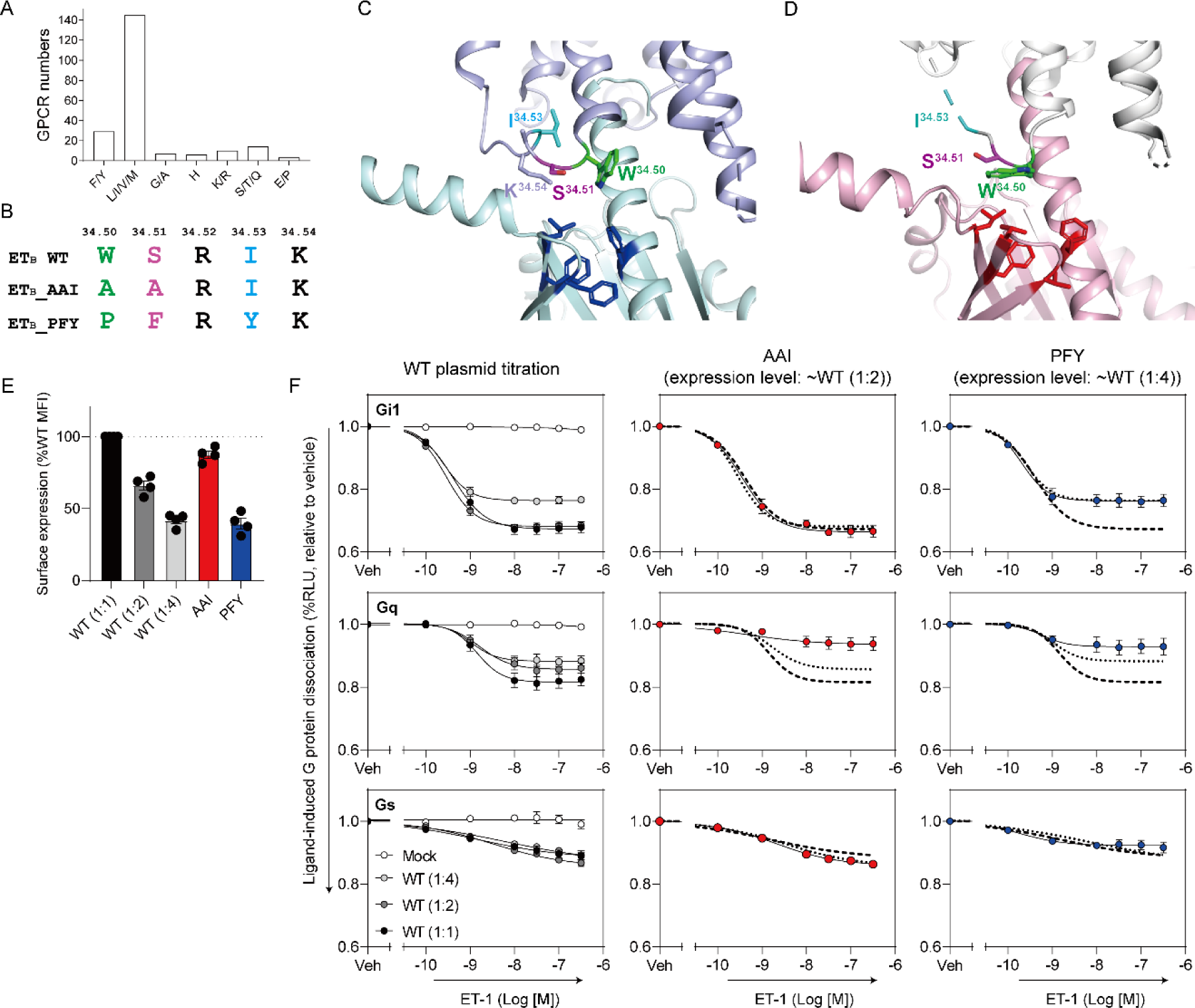
Role of ICL2 on ET_B_-mediated G protein activation. (A) The proportion of the residue 34.51 among the class A GPCRs. (B) Sequence alignment of ICL2 among ET_B_ wild-type (WT), ET_B_ AAI mutant, and ET_B_ PFY mutant. (C and D) The interaction of ET_B_ ICL2 with Gαi1 (C) and with Gαq (D). Selected residues are represented as sticks on the crystal structures of ET_B_-Gi1 (PDB: 8HBD) and ET_B_-Gq (PDB: 8HCX). The receptors are colored light purple and white, Gαi1 is colored light cyan, and Gαq is colored light pink. (E) Expression levels of the ICL2 mutant constructs. HEK293 cells expressing the indicated N-terminally FLAG-tagged ET_B_ constructs along with the NanoBiT-Gq sensor were subjected to the flow cytometer using an anti-FLAG-tag antibody. Symbols and error bars represent mean and standard error of the mean, respectively, derived from four independent experiments (dots), each conducted in duplicate. MFI, mean florescent intensity. The notation WT (1:1), WT (1:2), and WT (1:4) correspond to the volume ratios of transfected plasmids. (F) Effects of ET_B_ ICL2 mutation on ET_B_-mediated G protein activation. HEK293 cells expressing the indicated ET_B_ constructs along with the NanoBiT-G protein sensors were subjected to the NanoBiT-G protein assay. Symbols and error bars represent mean and standard error of the mean, respectively, derived from three independent experiments, each conducted in duplicate. Dashed lines in the AAI panels and the PFY panels denote WT (1:1) response curves. Dotted lines in the AAI panels and the PFY panels denote response curves in WT (1:2) and WT (1:4), respectively.

In addition to residue 34.51, the other residues in ICL2 comprise amino acids that are relatively well-conserved (Figure S2D–S2F); residue 34.50 is highly conserved as Pro, residue 34.52 is frequently conserved with positively charged amino acids (Lys or Arg), and residue 34.53 is frequently conserved with large hydrophobic amino acids (Phe, Tyr, or Trp).

Interestingly, however, ET_B_ has a unique composition of ICL2 residues (Figure 5B). ET_B_ possesses Ser at the residue 34.51, and therefore, there is no hydrophobic interaction between the residue 34.51 and hydrophobic pocket of Gαi1 or Gαq in the ET_B_-Gi1 or ET_B_-Gq structures (Figure 5C and 5D). Instead, Trp^34.50^ seemingly contributes to the hydrophobic interaction between ICL2 and Gαq (Figure 5D) but not Gαi1 (Figure 5C).

To investigate the role of ICL2 in ET_B_-induced G protein activation, we designed two ET_B_ mutants (Figure 5B) and measured G protein activation using the NanoBiT-G protein assay (Figure 5E-5F). In ET_B__AAI, the residues 34.50 and 34.51 were mutated to Ala. In ET_B__PFY, the residues 34.50, 34.51, and 34.53 were mutated to Pro, Phe, and Tyr, respectively, to introduce the most conserved amino acid residue in ICL2 (Figure S2D and S2F). In the Gi1 assay, we observed similar responses in ET_B__AAI and ET_B__PFY as compared with the expression level-matched wild-type (WT) (Figure 5F), showing that the residues 34.50, 34.51, and 34.53 are dispensable for ET_B_-induced Gi1 activation. This is consistent with the shallow interaction of the residue 34.51 with Gαi/o (Figure S2C) and barely any interaction of Gαi with Ser^34.51^ and Trp^34.50^ in the ET_B_-Gi1 structure (Figure 5C). Similarly, our previous report showed that the muscarinic acetylcholine receptor M2-induced GDP release from Gi was not much affected by Leu^34.51^ mutating to Ala^21^. In contrast, Gq activation was profoundly reduced in ET_B__AAI and ET_B__PFY compared to the expression level-matched WT (Figure 5F). The reduced Gq activation by ET_B__PFY was unexpected to us. We anticipated that switching the residue 34.51 into the conserved bulky hydrophobic amino acid would facilitate ET_B_-Gq coupling by forming a deep hydrophobic interaction with Gαq as in other GPCR-Gq complexes; however, this was not the case (Figure S2B)^32^. Therefore, it is tempting to suggest that ICL2 of ET_B_ is critical for activation of Gq but the structural mechanism behind this might be different from other GPCR-Gq coupling.

Finally, we tested whether alterations in the ICL2 sequence would affect ET_B_-Gs coupling. As described above, the bulky hydrophobic residue 34.51 of ICL2 makes an extensive contact with the hydrophobic pocket of Gαs (Figure S2A), and mutations of the residue 34.51 into another smaller residues reduce GPCR-Gs coupling or switched GPCR-Gs coupling to GPCR-Gi coupling^24,33–35^. Furthermore, introducing large hydrophobic amino acids into the residue 34.51 switched GPCR-Gi coupling to GPCR-Gs coupling^36^. Therefore, we hypothesized that ET_B__PFY would couple efficiently to Gs. However, neither ET_B__AAI nor ET_B__PFY showed increased ET_B_-Gs coupling (Figure 5F), suggesting that ICL2 of ET_B_ is not determining inefficient ET_B_-Gs coupling.

In summary, the unique sequence of ICL2 in ET_B_ is essential for ET_B_-Gq coupling but not for ET_B_-Gi1 coupling. Furthermore, the underlying structural mechanisms by which ICL2 contributes to the specificity of ETB-Gq coupling may exhibit differences when compared to the coupling mechanisms observed in other GPCR-G protein couplings.

## CONCLUSION

Herein, through the HDX analysis, we compared the conformational changes in full-length Gs, Gi1, and Gq upon binding to WT ET-1 bound ET_B_ without any auxiliary protein or modification to enhance stable complex formation. Our findings revealed varying degrees of coupling efficiency depending on G protein subtypes. Gi1 exhibited the most significant activation, followed by Gq, while Gs showed minimal activation. This observation challenges the conventional view of ET_B_ as a promiscuously coupled receptor and highlights the need for a deeper understanding of the molecular basis of GPCR-G protein interactions. By elucidating the structural determinants underlying the G protein selectivity exhibited by ET_B_, we provided valuable insights into the complexity of GPCR signaling pathways.

Of particular significance is the role of ICL2 in ET_B_ in modulating its selectivity towards different G protein subtypes. Our mutational analysis of key residues within ICL2 revealed that alterations in this region significantly affected ET_B_-Gq coupling but had minimal effects on ET_B_-Gi/o coupling. Of note, the orientations of G protein integration within the ET_B_-Gq configuration markedly deviated from those of the other GPCR-Gq complexes (Figure S3). Moreover, contrary to typical GPCR-Gq complexes, the interaction between the ICL2 of ET_B_ and hydrophobic pocket of Gαq is much small (Figure 5D). The cryoEM structures suggest that Trp^34.50^ of ET_B_ takes over the role of residue 34.51 in other GPCRs and maintains hydrophobic interactions. The unique composition of ICL2 in ET_B_ likely contributes to the unique orientation of the complex, thereby allowing for promiscuous G protein coupling. This is consistent with a recent study suggesting that the angle of ICL2 influences Gq-coupling^37^.

Despite extensive efforts, we were unable to identify binding interfaces between Gi1/Gq and ET_B_ using our current HDX-MS analysis. What was more unexpected is that the C-terminus of Gα α5 was not affected by ET_B_ co-incubation in ET_B_-Gi and ET_B_-Gq HDX-MS analyses. These data imply that the interaction between ET_B_ and Gi1/Gq is unstable probably due to the unstable and/or shallow interaction of the C-terminus of Gα α5 to ET_B_. Despite this unstable and/or shallow interaction of the C-terminus of Gα α5 to ET_B_, the C-terminus of Gα α5 was still critical for ET_B_-Gi and ET_B_-Gq coupling. We speculate that the intriguing interplay between shallow interactions and critical coupling suggests a potential contributory role to the promiscuity exhibited by ET_B_.

In conclusion, the interaction of the C-terminus of Gα α5 to ET_B_ is unstable and/or shallow but critical for ET_B_-Gi and ET_B_-Gq coupling while the interaction of ICL2 is critical only for ET_B_-Gq coupling but not for ET_B_-Gi coupling. This suggests a nuanced interplay between the ET_B_ and its associated G proteins, wherein distinct structural motifs within the ET_B_ contribute to the differential coupling efficiencies toward G protein subtypes. However, the specific interactions influencing ET_B_-G protein coupling have yet to be identified; further investigation is needed to comprehensively understand the molecular mechanisms governing ET_B_-mediated signaling.

## SUPPLEMENTAL INFORMATION

Supplemental information includes three figures and one supplementary dataset and can be found online.

## ACKNOWLEDGEMENTS

We thank Kayo Sato, Shigeko Nakano, and Ayumi Inoue at Tohoku University for their assistance with plasmid preparation and the cell-based GPCR assays. This work was supported by grants from the National Research Foundation of Korea funded by the Korean government (NRF-2021R1A2C3003518 and NRF-2019R1A5A2027340 to K.Y.C.) and by a grant from the Ministry of Oceans and Fisheries’ R&D project, Korea (2021633). W.S. was funded by KAKENHI 22H02751. O.N. was funded by KAKENHI 21H05037. A.I. was funded by KAKENHI JPW21H04791, JP21H05113, JP21H05037, and JPJSBP120218801 from Japan by the Society for the Promotion of Science (JSPS); JPMJFR215T, JPMJMS2023, and 22714181 from the Japan Science and Technology Agency (JST); and JP22ama121038 and JP22zf0127007 from the Japan Agency for Medical Research and Development (AMED).

## AUTHOR CONTRIBUTIONS

D.H. prepared Gs, Gi1, and Gq, and performed HDX-MS for all samples. W.S. prepared ET_B_ with the supervision by O.N. A.I. generated mutant constructs and performed the NanoBiT assays and the flow cytometry analysis. K.Y.C. and A.I. initiated and supervised the project. K.Y.C., D.H., and A.I. analyzed the data and wrote the manuscript.

## DECLARATION OF INTERESTS

O.N. is a co-founder and scientific advisor for Curreio. All other authors declare no competing interests.

## INCLUSION AND DIVERSITY

We support inclusive, diverse, and equitable conduct of research.

## METHODS

### RESOURCE AVAILABILITY

#### Lead contact

Further information and requests for reagents should be directed to and fulfilled by the lead contact, Ka Young Chung.

#### Materials availability

All unique/stable reagents generated in this study are available from the lead contact with a complete materials transfer agreement.

#### Data and code availability

Data reported in this paper will be shared by the lead contact upon request.

### EXPERIMENTAL MODEL AND SUBJECT DETAILS

Human Gαs was expressed in *Escherichia coli* BL21(DE3) strain, and human Gαq, human Gαi1, human His6-Gβ_1_, bovine Gγ_2_, and human ETB were expressed in the sf9 cell-line from *Spodoptera frugiperda* pupal ovarian tissue.

## METHODS DETAILS

### Protein expression and purification

WT human Gαq, human Gαi1, human His6-Gβ_1_, and bovine Gγ_2_ were cloned into the pVL1392 vector, and Ric8A was cloned into the pFastBac1 vector. WT human Gαs containing an N-terminal 6XHis-tag and a TEV cleavage site was cloned into the pET28a vector. Heterotrimeric Gq and Gi1 were expressed and purified as previously established^21^. For heterotrimeric Gs, Gαs and Gβ_1_γ_2_ were expressed and purified respectively as previously described^24^. The full-length human ET_B_ gene was subcloned into a pFastBac vector with an HA signal peptide sequence at the N-terminus. A FLAG epitope tag (DYKDDDK) was introduced between residues G57 and L66. The native signal peptide was replaced with the haemagglutinin signal peptide. ET-1-bound WT ET_B_ was purified in dodecyl maltoside (DDM) and cholesteryl hemisuccinate (CHS), as previously established^38^.

### Co-incubation protocol

To form the ET_B_-Gq, ET_B_-Gi1, and ET_B_-Gs complexes, ET-1-bound ET_B_ and heterotrimeric G proteins were mixed at a final concentration of 50 μM at room temperature for 4 h. Apyrase (200 mU/mL) was added after 90 min of incubation to hydrolyze released GDP.

### HDX-MS

For the ET_B_-G protein complexes, hydrogen/deuterium exchange was initiated by mixing 5 μL of protein sample and 25 μL of D_2_O buffer (20 mM HEPES, pD 7.4, 100 mM NaCl, 0.1% DDM, 1 mM MgCl_2_, and 100 μM TCEP supplemented with 5 μM ET-1 or 10 μM GDP for the complex or alone samples, respectively) and incubated for 10, 100, 1,000, and 10,000 s at room temperature. The deuterated samples were quenched using 30 μL of ice-cold quench buffer (60 mM NaH_2_PO_4_, pH 2.01, 20 mM TCEP, and 10% glycerol), snap-frozen on dry ice, and stored at -80 °C. Non-deuterated (ND) samples were prepared by mixing 5 μL of the protein sample with 25 μL of their respective H_2_O buffers, followed by quenching and freezing, as described above.

The quenched samples were digested by passing through an immobilized pepsin column (2.1 × 30 mm; Life Technologies, Carlsbad, CA, USA) at a flow rate of 100 µL/min with 0.05% formic acid in H_2_O at 12 °C. The peptide fragments were subsequently collected on a C18 VanGuard trap column (1.7 µm × 30 mm; Waters, Milford, MA, USA) and desalted with 0.05% formic acid in H_2_O. Peptic peptides were then separated using ultra-pressure liquid chromatography (UPLC) on an ACQUITY UPLC C18 column (1.7 µm, 1.0 mm × 100 mm; Waters) at 40 µL/min with an acetonitrile gradient created by two pumps— mobile phase A (0.15% formic acid in H_2_O) and B (0.15% formic acid in acetonitrile). The gradient started at 8% B and increased to 85% B over 8.5 min. To minimize the back-exchange of deuterium to hydrogen, the sample, solvents, trap, and UPLC column were all maintained at pH 2.5 and 0.5 °C during analysis. Mass spectrometry analyses were performed using a Xevo G2-XS QTof (Waters) as established previously^21,24^. Peptides were identified in the ND samples using the ProteinLynx Global Server 2.4 (Waters). To process the HDX-MS data, the amount of deuterium in each peptide was determined by measuring the centroid of the isotopic distribution using DynamX 3.0 (Waters).

### NanoBiT G protein dissociation assay

Ligand-induced G protein dissociation was measured using a NanoBiT-G-protein dissociation assay, whereby the interaction between a Gα and Gβγ subunits which were transiently expressed in HEK293 cells was monitored by the NanoBiT system as established previously^19^. The ET_B_ constructs used in the cell-based assays contained an N-terminal FLAG epitope tag and was described previously^39^.

### Flow cytometry analysis

Cell-surface expression levels of ET_B_ mutant constructs were quantified by flow cytometry analysis as described previously^40^. Briefly, HEK293 cells transiently expressing ET_B_ construct along with the NanoBiT-Gq sensor were fluorescently labeled with anti-FLAG epitope tag primary antibody (Clone 1E6, FujiFilm Wako Pure Chemicals) and Alexa 488-conjugated secondary antibody (Thermo Fisher Scientific). Fluorescent counts from individual cells were measured by a flow cytometer (EC800, Sony). For each experiment, we normalized an MFI value of the mutants by that of WT performed in parallel and denoted relative levels.

## QUANTIFICATION AND STATISTICAL ANALYSIS

For HDX-MS analysis, the Student’s *t*-test was used to assess the statistically significant differences between samples with and without the binding partner. One-way analysis of variance, followed by Tukey’s post-hoc test, was used to analyze the differences between more than three conditions. Statistical significance was at p <0.05. More than three independent experiments were performed for each dataset.

## REFERENCES

1. Decker, E.R., and Brock, T.A. (1998). Endothelin and Calcium Signaling. In Endothelin Receptors and Signaling Mechanisms, D.M. Pollock, and R.F. Highsmith, eds. (Springer Berlin Heidelberg), pp. 131–146. 10.1007/978-3-662-11672-2_10.

2. Arai, H., Hori, S., Aramori, I., Ohkubo, H., and Nakanishi, S. (1990). Cloning and expression of a cDNA encoding an endothelin receptor. Nature 348, 730–732. 10.1038/348730a0.

3. Sakurai, T., Yanagisawa, M., Takuwa, Y., Miyazaki, H., Kimura, S., Goto, K., and Masaki, T. (1990). Cloning of a cDNA encoding a non-isopeptide-selective subtype of the endothelin receptor. Nature 348, 732–735. 10.1038/348732a0.

4. Sokolovsky, M. (1995). Endothelin receptor subtypes and their role in transmembrane signaling mechanisms. Pharmacol Ther 68, 435–471. 10.1016/0163-7258(95)02015-2.

5. Takigawa, M., Sakurai, T., Kasuya, Y., Abe, Y., Masaki, T., and Goto, K. (1995). Molecular identification of guanine-nucleotide-binding regulatory proteins which couple to endothelin receptors. Eur J Biochem 228, 102–108. 10.1111/j.1432-1033.1995.tb20236.x.

6. Okashah, N., Wan, Q., Ghosh, S., Sandhu, M., Inoue, A., Vaidehi, N., and Lambert, N.A. (2019). Variable G protein determinants of GPCR coupling selectivity. Proc Natl Acad Sci U S A 116, 12054–12059. 10.1073/pnas.1905993116.

7. Inoue, A., Raimondi, F., Kadji, F.M.N., Singh, G., Kishi, T., Uwamizu, A., Ono, Y., Shinjo, Y., Ishida, S., Arang, N., et al. (2019). Illuminating G-Protein-Coupling Selectivity of GPCRs. Cell 177, 1933–1947 e1925. 10.1016/j.cell.2019.04.044.

8. Avet, C., Mancini, A., Breton, B., Le Gouill, C., Hauser, A.S., Normand, C., Kobayashi, H., Gross, F., Hogue, M., Lukasheva, V., et al. (2022). Effector membrane translocation biosensors reveal G protein and betaarrestin coupling profiles of 100 therapeutically relevant GPCRs. Elife 11. 10.7554/eLife.74101.

9. Masuho, I., Kise, R., Gainza, P., Von Moo, E., Li, X., Tany, R., Wakasugi-Masuho, H., Correia, B.E., and Martemyanov, K.A. (2023). Rules and mechanisms governing G protein coupling selectivity of GPCRs. Cell Reports 42, 113173. 10.1016/j.celrep.2023.113173.

10. Harding, S.D., Armstrong, J.F., Faccenda, E., Southan, C., Alexander, S.P.H., Davenport, A.P., Spedding, M., and Davies, J.A. (2023). The IUPHAR/BPS Guide to PHARMACOLOGY in 2024. Nucleic Acids Res 10.1093/nar/gkad944.

11. Hauser, A.S., Avet, C., Normand, C., Mancini, A., Inoue, A., Bouvier, M., and Gloriam, D.E. (2022). Common coupling map advances GPCR-G protein selectivity. eLife 11, e74107. 10.7554/eLife.74107.

12. Sano, F.K., Akasaka, H., Shihoya, W., and Nureki, O. (2023). Cryo-EM structure of the endothelin-1-ET(B)-G(i) complex. Elife 12. 10.7554/eLife.85821.

13. Ji, Y., Duan, J., Yuan, Q., He, X., Yang, G., Zhu, S., Wu, K., Hu, W., Gao, T., Cheng, X., et al. (2023). Structural basis of peptide recognition and activation of endothelin receptors. Nat Commun 14, 1268 10.1038/s41467-023-36998-9.

14. Shihoya, W., Sano, F.K., and Nureki, O. (2023). Structural insights into endothelin receptor signalling. J Biochem 174, 317–325. 10.1093/jb/mvad055.

15. Dixon, A.S., Schwinn, M.K., Hall, M.P., Zimmerman, K., Otto, P., Lubben, T.H., Butler, B.L., Binkowski, B.F., Machleidt, T., Kirkland, T.A., et al. (2016). NanoLuc Complementation Reporter Optimized for Accurate Measurement of Protein Interactions in Cells. ACS Chem Biol 11, 400–408. 10.1021/acschembio.5b00753.

16. Hvidt, A., and Nielsen, S.O. (1966). Hydrogen Exchange in Proteins. In Advances in Protein Chemistry, C.B. Anfinsen, M.L. Anson, J.T. Edsall, and F.M. Richards, eds. (Academic Press), pp. 287–386. 10.1016/S0065-3233(08)60129-1.

17. Wales, T.E., and Engen, J.R. (2006). Hydrogen exchange mass spectrometry for the analysis of protein dynamics. Mass Spectrom Rev 25, 158–170. 10.1002/mas.20064.

18. Qu, C., Park, J.Y., Yun, M.W., He, Q.T., Yang, F., Kim, K., Ham, D., Li, R.R., Iverson, T.M., Gurevich, V.V., et al. (2021). Scaffolding mechanism of arrestin-2 in the cRaf/MEK1/ERK signaling cascade. Proc Natl Acad Sci U S A 118. 10.1073/pnas.2026491118.

19. Ham, D., Inoue, A., Xu, J., Du, Y., and Chung, K.Y. (2024). Molecular mechanism of muscarinic acetylcholine receptor M3 interaction with Gq. Commun Biol 7, 362. 10.1038/s42003-024-06056-1.

20. Chung, K.Y., Rasmussen, S.G., Liu, T., Li, S., DeVree, B.T., Chae, P.S., Calinski, D., Kobilka, B.K., Woods, V.L., Jr., and Sunahara, R.K. (2011). Conformational changes in the G protein Gs induced by the beta2 adrenergic receptor. Nature 477, 611–615. 10.1038/nature10488.

21. Kim, H.R., Xu, J., Maeda, S., Duc, N.M., Ahn, D., Du, Y., and Chung, K.Y. (2020). Structural mechanism underlying primary and secondary coupling between GPCRs and the Gi/o family. Nat Commun 11, 3160. 10.1038/s41467-020-16975-2.

22. Rasmussen, S.G.F., DeVree, B.T., Zou, Y., Kruse, A.C., Chung, K.Y., Kobilka, T.S., Thian, F.S., Chae, P.S., Pardon, E., Calinski, D., et al. (2011). Crystal structure of the β2 adrenergic receptor–Gs protein complex. Nature 477, 549–555. 10.1038/nature10361.

23. Ahn, D., Provasi, D., Duc, N.M., Xu, J., Salas-Estrada, L., Spasic, A., Yun, M.W., Kang, J., Gim, D., Lee, J., et al. (2023). Gαs slow conformational transition upon GTP binding and a novel Gαs regulator. iScience 26. 10.1016/j.isci.2023.106603.

24. Du, Y., Duc, N.M., Rasmussen, S.G.F., Hilger, D., Kubiak, X., Wang, L., Bohon, J., Kim, H.R., Wegrecki, M., Asuru, A., et al. (2019). Assembly of a GPCR-G Protein Complex. Cell 177, 1232–1242 e1211. 10.1016/j.cell.2019.04.022.

25. Lee, Y., Warne, T., Nehmé, R., Pandey, S., Dwivedi-Agnihotri, H., Chaturvedi, M., Edwards, P.C., García-Nafría, J., Leslie, A.G.W., Shukla, A.K., and Tate, C.G. (2020). Molecular basis of β-arrestin coupling to formoterol-bound β(1)-adrenoceptor. Nature 583, 862–866. 10.1038/s41586-020-2419-1.

26. Flock, T., Hauser, A.S., Lund, N., Gloriam, D.E., Balaji, S., and Babu, M.M. (2017). Selectivity determinants of GPCR-G-protein binding. Nature 545, 317–322. 10.1038/nature22070.

27. Hilger, D., Masureel, M., and Kobilka, B.K. (2018). Structure and dynamics of GPCR signaling complexes. Nat Struct Mol Biol 25, 4–12. 10.1038/s41594-017-0011-7.

28. Glukhova, A., Draper-Joyce, C.J., Sunahara, R.K., Christopoulos, A., Wootten, D., and Sexton, P.M. (2018). Rules of Engagement: GPCRs and G Proteins. ACS Pharmacol Transl Sci 1, 73–83. 10.1021/acsptsci.8b00026.

29. Ballesteros, J.A., and Weinstein, H. (1995). [19] Integrated methods for the construction of three-dimensional models and computational probing of structure-function relations in G protein-coupled receptors. In Receptor Molecular Biology, S.C. Sealfon, ed. (Academic Press), pp. 366–428. 10.1016/s1043-9471(05)80049-7.

30. Koehl, A., Hu, H., Maeda, S., Zhang, Y., Qu, Q., Paggi, J.M., Latorraca, N.R., Hilger, D., Dawson, R., Matile, H., et al. (2018). Structure of the micro-opioid receptor-G(i) protein complex. Nature 558, 547–552. 10.1038/s41586-018-0219-7.

31. Moro, O., Lameh, J., Högger, P., and Sadée, W. (1993). Hydrophobic amino acid in the i2 loop plays a key role in receptor-G protein coupling. Journal of Biological Chemistry 268, 22273–22276. 10.1016/s0021-9258(18)41524-4.

32. Maeda, S., Qu, Q., Robertson, M.J., Skiniotis, G., and Kobilka, B.K. (2019). Structures of the M1 and M2 muscarinic acetylcholine receptor/G-protein complexes. Science 364, 552–557. 10.1126/science.aaw5188.

33. Chen, X.P., Yang, W., Fan, Y., Luo, J.S., Hong, K., Wang, Z., Yan, J.F., Chen, X., Lu, J.X., Benovic, J.L., and Zhou, N.M. (2010). Structural determinants in the second intracellular loop of the human cannabinoid CB1 receptor mediate selective coupling to G(s) and G(i). Br J Pharmacol 161, 1817–1834. 10.1111/j.1476-5381.2010.01006.x.

34. Suno, R., Sugita, Y., Morimoto, K., Takazaki, H., Tsujimoto, H., Hirose, M., Suno-Ikeda, C., Nomura, N., Hino, T., Inoue, A., et al. (2022). Structural insights into the G protein selectivity revealed by the human EP3-G(i) signaling complex. Cell Rep 40, 111323. 10.1016/j.celrep.2022.111323.

35. Zhao, L.H., Lin, J., Ji, S.Y., Zhou, X.E., Mao, C., Shen, D.D., He, X., Xiao, P., Sun, J., Melcher, K., et al. (2022). Structure insights into selective coupling of G protein subtypes by a class B G protein-coupled receptor. Nat Commun 13, 6670. 10.1038/s41467-022-33851-3.

36. Zheng, C., Chen, L., Chen, X., He, X., Yang, J., Shi, Y., and Zhou, N. (2013). The second intracellular loop of the human cannabinoid CB2 receptor governs G protein coupling in coordination with the carboxyl terminal domain. PloS one 8, e63262. 10.1371/journal.pone.0063262.

37. Powers, A.S., Khan, A., Paggi, J.M., Latorraca, N.R., Souza, S., Salvo, J.D., Lu, J., Soisson, S.M., Johnston, J.M., Weinglass, A.B., and Dror, R.O. (2023). A non-canonical mechanism of GPCR activation. bioRxiv : the preprint server for biology. 10.1101/2023.08.14.553154.

38. Shihoya, W., Nishizawa, T., Okuta, A., Tani, K., Dohmae, N., Fujiyoshi, Y., Nureki, O., and Doi, T. (2016). Activation mechanism of endothelin ET(B) receptor by endothelin-1. Nature 537, 363–368. 10.1038/nature19319.

39. Shihoya, W., Nishizawa, T., Yamashita, K., Inoue, A., Hirata, K., Kadji, F.M.N., Okuta, A., Tani, K., Aoki, J., Fujiyoshi, Y., et al. (2017). X-ray structures of endothelin ET(B) receptor bound to clinical antagonist bosentan and its analog. Nat Struct Mol Biol 24, 758–764. 10.1038/nsmb.3450.

40. Nagiri, C., Shihoya, W., Inoue, A., Kadji, F.M.N., Aoki, J., and Nureki, O. (2019). Crystal structure of human endothelin ET(B) receptor in complex with peptide inverse agonist IRL2500. Commun Biol 2, 236. 10.1038/s42003-019-0482-7.

