## Supplementary figures for "Molecular mechanism of the endothelin receptor type B interactions with Gs, Gi, and Gq"

### Slide 1
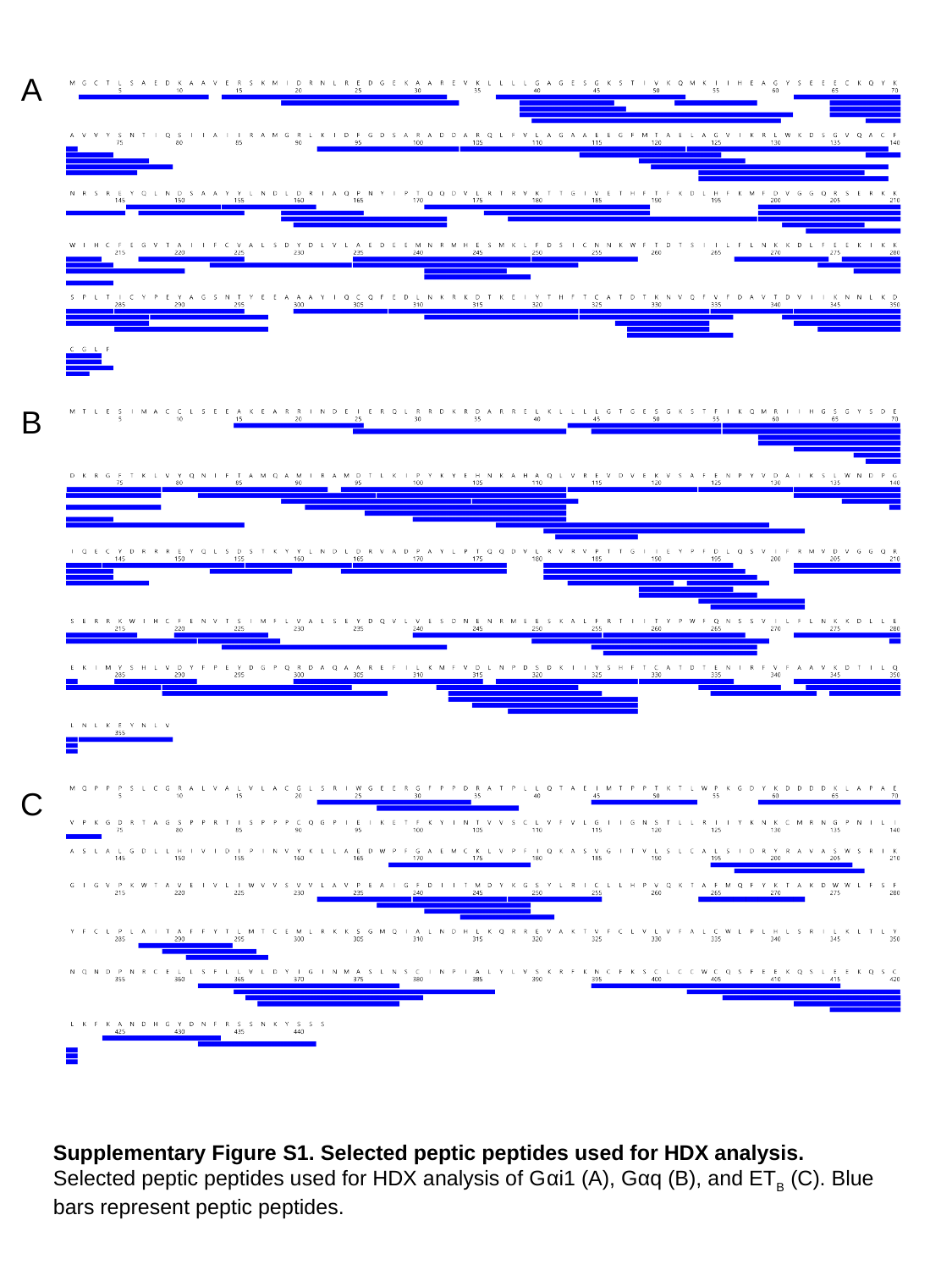

A
B
C
Supplementary Figure S1. Selected peptic peptides used for HDX analysis.
Selected peptic peptides used for HDX analysis of Gαi1 (A), Gαq (B), and ET_B_ (C). Blue bars represent peptic peptides.

### Slide 2
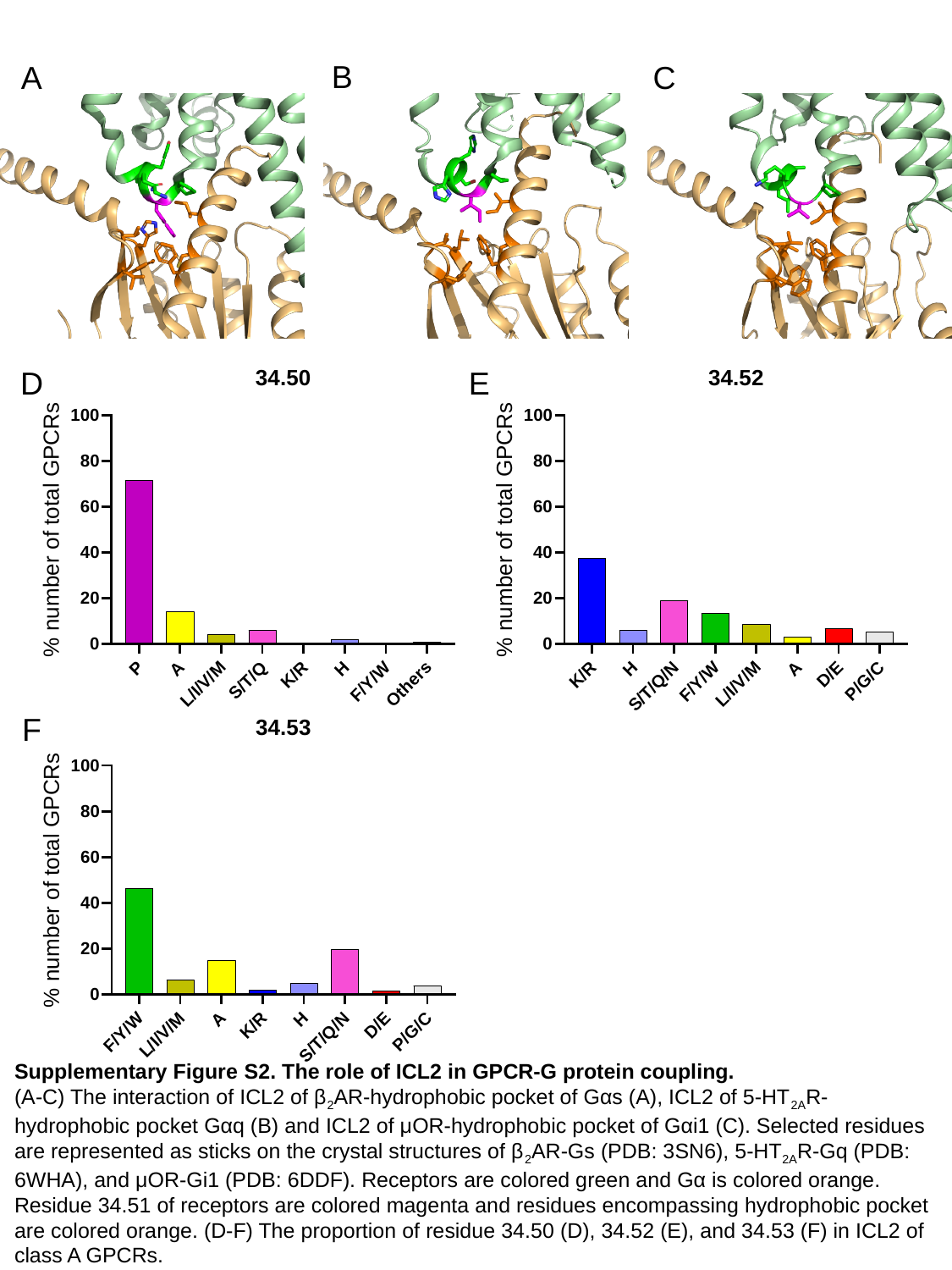

B
A
C
D
E
F
Supplementary Figure S2. The role of ICL2 in GPCR-G protein coupling.
(A-C) The interaction of ICL2 of β_2_AR-hydrophobic pocket of Gαs (A), ICL2 of 5-HT_2A_R-hydrophobic pocket Gαq (B) and ICL2 of μOR-hydrophobic pocket of Gαi1 (C). Selected residues are represented as sticks on the crystal structures of β_2_AR-Gs (PDB: 3SN6), 5-HT_2A_R-Gq (PDB: 6WHA), and μOR-Gi1 (PDB: 6DDF). Receptors are colored green and Gα is colored orange. Residue 34.51 of receptors are colored magenta and residues encompassing hydrophobic pocket are colored orange. (D-F) The proportion of residue 34.50 (D), 34.52 (E), and 34.53 (F) in ICL2 of class A GPCRs.

### Slide 3
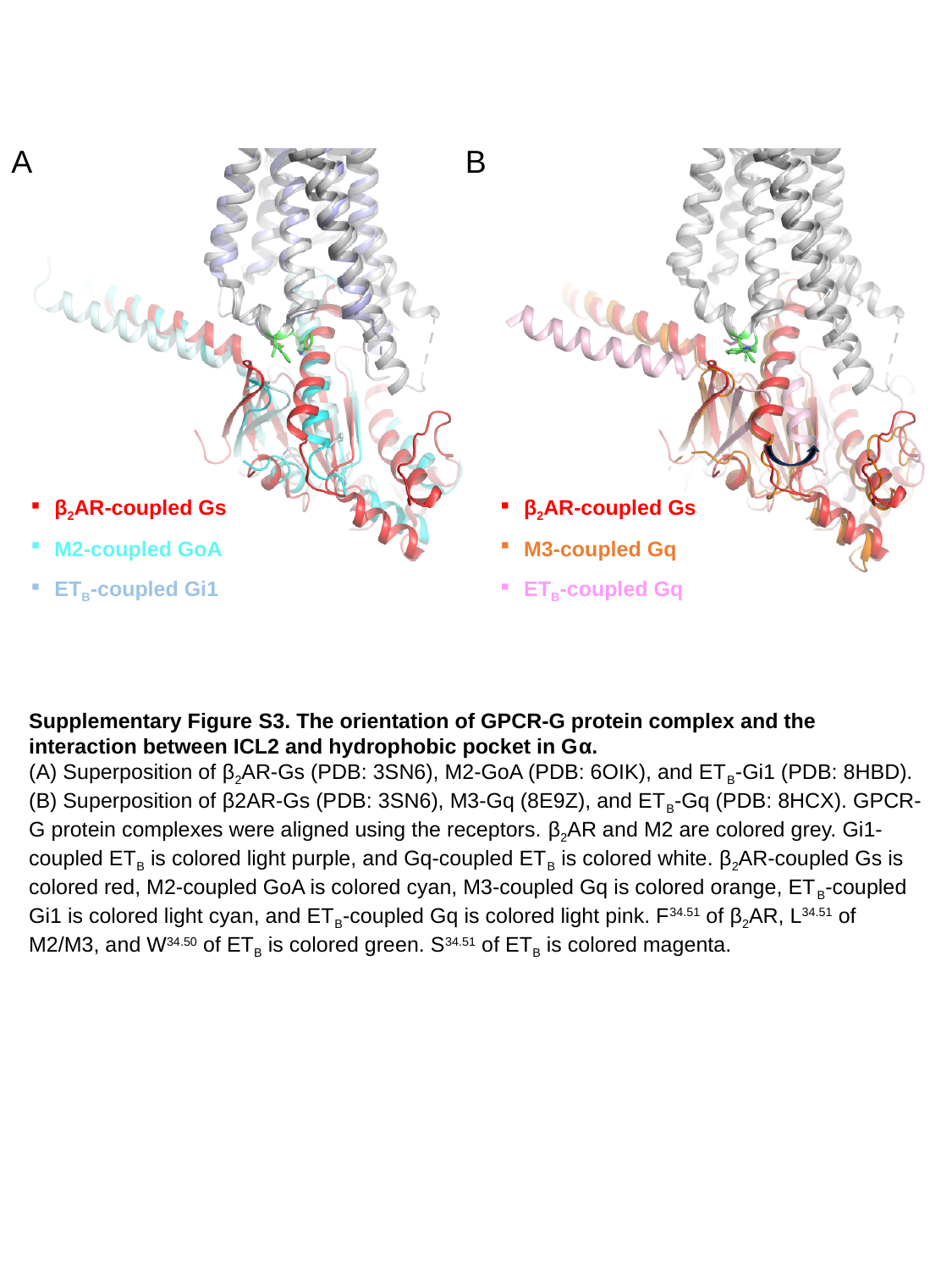

A
B
β_2_AR-coupled Gs
M2-coupled GoA
ET_B_-coupled Gi1
β_2_AR-coupled Gs
M3-coupled Gq
ET_B_-coupled Gq
Supplementary Figure S3. The orientation of GPCR-G protein complex and the interaction between ICL2 and hydrophobic pocket in Gα.
(A) Superposition of β_2_AR-Gs (PDB: 3SN6), M2-GoA (PDB: 6OIK), and ET_B_-Gi1 (PDB: 8HBD). (B) Superposition of β2AR-Gs (PDB: 3SN6), M3-Gq (8E9Z), and ET_B_-Gq (PDB: 8HCX). GPCR-G protein complexes were aligned using the receptors. β_2_AR and M2 are colored grey. Gi1-coupled ET_B_ is colored light purple, and Gq-coupled ET_B_ is colored white. β_2_AR-coupled Gs is colored red, M2-coupled GoA is colored cyan, M3-coupled Gq is colored orange, ET_B_-coupled Gi1 is colored light cyan, and ET_B_-coupled Gq is colored light pink. F^34.51^ of β_2_AR, L^34.51^ of M2/M3, and W^34.50^ of ET_B_ is colored green. S^34.51^ of ET_B_ is colored magenta.
